# Modeling Complex Effects and Individual Variability in Multi-Paradigm fMRI with Nonlinear Mixed Models

**DOI:** 10.64898/2026.05.16.725673

**Authors:** Xiaoxuan Li, Gemeng Zhang, Gang Qu, Anton Orlichenko, Zhengming Ding, Tony W Wilson, Julia M Stephen, Vince D Calhoun, Yu-Ping Wang

**Author notes:** (Corresponding author: Yu-Ping Wang).

## Abstract

Functional magnetic resonance imaging (fMRI) data are inherently complex, characterized by high dimensionality, intricate inter-regional dependencies, and substantial individual variability across experimental paradigms. Traditional linear mixed models (LMMs) provide a principled framework that models population-level fixed effects while estimating variance components arising from subject-level random effects; however, they often fail to adequately capture nonlinear relationships inherent in neuroimaging data. To address these limitations, we introduce the nonlinear mixed model (NMM) approach, an innovative extension of the LMM framework that integrates neural networks to flexibly model complex fixed-effect relationships while preserving the random-effects structure to account for individual differences. NMM advances fMRI analysis by: (1) identifying robust functional connectivity (FC) patterns consistently observed across multiple paradigms; (2) leveraging SHapley Additive exPlanations (SHAP) analysis to provide post-hoc interpretability of the nonlinear fixed effects, quantifying how age, sex, and paradigm contribute to predicted FC and how these effects are distributed across large-scale brain networks; and (3) using subject-specific random effects as neural fingerprints that not only show systematic variability across attention and default mode systems but also predict standardized cognitive scores, demonstrating biological relevance. Applied to the Philadelphia Neurodevelopmental Cohort (PNC) across emotion, n-back, and resting-state paradigms, NMM achieved superior model fit relative to classical LMMs, as evidenced by lower mean squared error (MSE) in predicting FC. This framework offers a statistically rigorous and practically explainable approach for modeling large-scale FC from modest covariates while explicitly separating population-level effects from stable individual variability in functional brain organization.

## I Introduction

**T**HE past several decades have witnessed a paradigm shift in neuroscience from regional analyses to whole-rain network investigations, driven largely by advances in neuroimaging methods [1]–[5]. A key concept in this systems-level approach is functional connectivity (FC), defined as the statistical dependence between time series of neural activity originating from different brain regions [6]. Typically measured using correlation of blood-oxygen-level-dependent (BOLD) signals in functional magnetic resonance imaging (fMRI), FC has emerged as a fundamental metric for characterizing brain organization [7]. FC has been widely used to reveal organizational principles of brain function in both typical development and clinical conditions [8]–[12].

A critical step in FC studies is identifying between-group differences, which traditionally relies on statistical methods such as t-tests, ANOVA, or linear regression [13], [14]. While efficient, these conventional approaches based on the general linear model often struggle to model the complex interdependencies between brain regions inherent in neuroimaging data. This limitation is particularly relevant given the well established finding that FC exhibits substantial individual variability, serving as a unique “fingerprint” that can differentiate subjects across different scanning paradigms [15]–[18]. To address this, the Linear Mixed Model (LMM) has been introduced to fMRI group analysis, offering a robust framework for partitioning variance into group level fixed effects and individual specific random effects [19]. Narayan *et al*. [20] introduced a resampling-based mixed-effects framework for multi-subject FCs that models within-subject resamples and between-subject random effects, improving covariate inference. Results from the Adolescent Brain Cognitive Development (ABCD) study show that LMMs provide a principled framework for modeling complex neuroimaging data with multi-level dependencies such as site, family, and longitudinal measures, facilitating valid inference at the population level [21]. Moreover, whole-brain FC is typically represented as a connectivity matrix with tens of thousands of pairwise connections when atlases with hundreds of regions are used, and this rich structure is often not fully captured by simple linear models.

Concurrently, there is a growing recognition on the value of multi-paradigm fMRI data [22]–[25]. While the majority of FC research has focused on a single modality, typically resting-state fMRI [26], [27], studies increasingly acquire data under multiple task conditions (e.g., emotion, working memory) to provide complementary information about brain function. It has been shown that task-based FC can sometimes yield superior predictive power for behavioral and cognitive traits compared to resting-state FC, as tasks engage specific brain circuits relevant to the traits of interest [28], [29]. This suggests that integrating multiple fMRI paradigms within a unified analytical framework can harness their synergistic potential, leading to a more comprehensive understanding of brain-behavior relationships. Despite notable efforts in multi-task learning models that jointly analyze features from multiple modalities, these approaches often overlook the crucial aspect of inter-subject variability captured by the individual’s unique “fingerprint” across paradigms.

In this paper, we bridge two important methodological directions by proposing a novel framework that integrates the power of deep learning with the statistical rigor of LMMs for the analysis of multi-paradigm fMRI data. Specifically, we introduce the **Noninear Mixed Model (NMM)**, a hybrid approach for modeling FC, in which fixed effects are represented by a flexible neural network, and individual variability is explicitly modeled through random effects.

We evaluate **NMM** on data from the Philadelphia Neurodevelopmental Cohort (PNC), utilizing FC derived from three paradigms: emotion identification, n-back working memory, and resting state. Beyond achieving superior odel fit, a primary objective of our framework is to isolate subject-specific random effects (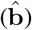). We hypothesized that by using the neural network to mathematically strip away the variance induced by age, sex, and task paradigm, the resulting random effects would encapsulate a pure, intrinsic representation of the individual’s functional architecture. Consequently, these extracted parameters should serve as highly effective predictors for general intelligence and cognitive capacities.

Furthermore, we integrate SHapley Additive exPlanations (SHAP) analysis [30], [31] to interpret the fixed effects both globally and across functional brain networks. Our results show that incorporating neural networks for fixed effects, coupled with explicit modeling of individual variance, not only enhances predictive performance but also yields random-effect estimates that reflect meaningful cognitive-behavioral associations. The overall architecture of our proposed approach is illustrated in Figure 1. To our knowledge, this is among the first studies to integrate an LMM framework with neural networks for modeling the relationship between FC and demographic as well as task covariates, explicitly accounting for individual variability while leveraging the rich information provided by multi-paradigm fMRI data.

**Fig. 1.**
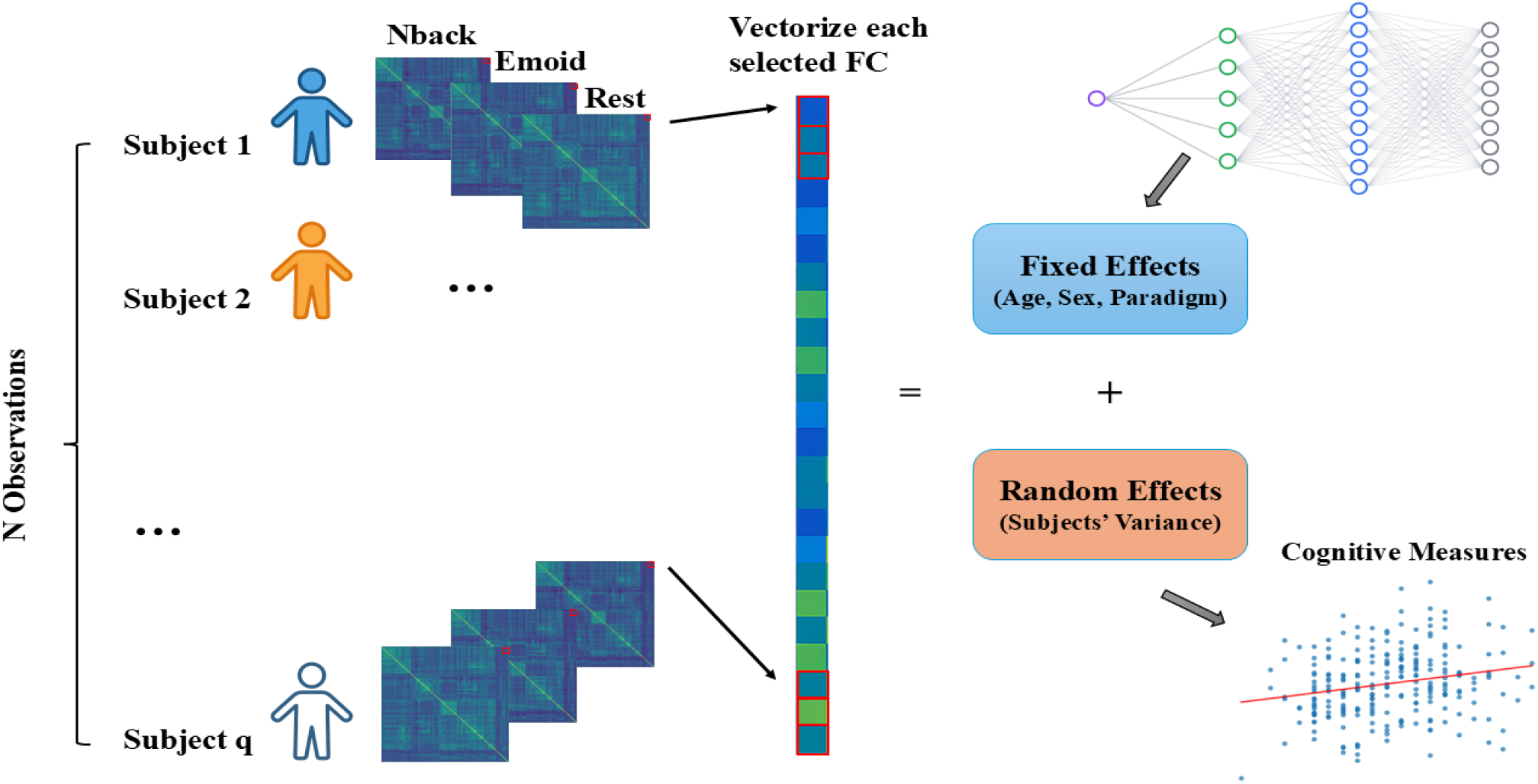
An overview of the analytical pipeline is presented. First, whole-brain FC matrices were derived for each subject and paradigm. Second, each selected pairwise FC value served as the dependent variable in an independent regression model. The fixed effects in these models were constructed via a MLP, which incorporated age, sex, and paradigm as input features. Third, model parameters estimated across all FC pairs were integrated to map whole-brain connectivity effects. To account for subject-level random effects, follow-up analyses were conducted to examine associations with cognitive measures.

## II. Dataset and Methods

### A. Data Acquisition

In this study, we utilized the PNC dataset, a large-scale publicly available resource developed through a collaboration between the Brain Behavior Laboratory at the University of Pennsylvania and the Center for Applied Genomics at the Children’s Hospital of Philadelphia [32]. The PNC was designed to characterize the interactions among brain imaging, behavior, and genetics in a diverse sample of adolescents [33]. A total of approximately 1445 participants aged 8–23 years underwent multi-modal neuroimaging, cognitive testing, and genotyping. All data are accessible via the dbGaP repository (https://dbgap.ncbi.nlm.nih.gov/home).

We selected a subset of participants who completed three fMRI paradigms: two task-based sessions—the n-back working memory task (nback) and the emotion identification task (emotion)—and one resting-state session. During the nback-fMRI, participants viewed a series of fractal images and were instructed to respond when the current stimulus matched the one presented two steps earlier. In the emotion-fMRI task, participants identified emotional expressions (e.g., happy, sad, angry, fearful, or neutral) from presented faces. Both tasks are well-validated probes of executive function and emotional perception and have previously been associated with intelligence-related measures [29], [34]–[36].

General cognitive ability was assessed using the Wide Range Achievement Test (WRAT) total score, derived from a computerized neurocognitive battery (CNB) [37]. The WRAT assesses core academic skills, including reading recognition, spelling, and arithmetic computation, and serves as a reliable proxy for Intelligence Quotient (IQ). Fluid intelligence was further evaluated using the Penn Matrix Reasoning Test (PMAT) and the Penn Verbal Reasoning Test (PVRT). After excluding participants with missing imaging data or clinical scores, a final sample of 1238 subjects (age 8–23 years, mean = 15.42; WRAT score 67–145, mean = 102.47; PMAT score 1.00–24.00, mean = 11.86; PVRT score 0.00–15.00, mean = 11.11; 668 females, 570 males) was retained for the main analyses. See details in Table 1.

**TABLE 1.** Demographics and clinical characteristics of the pnc dataset

| Variable | N or Mean ± SD |
| --- | --- |
| Sample size | $N = 1,238$ |
| Age (years) | $15.42 \pm 3.50$ |
| Sex (Female / Male) | 668/570 |
| WRAT | $102.47 \pm 16.07$ |
| PMAT | $11.86 \pm 4.10$ |
| PVRT | $11.11 \pm 2.87$ |

Note: Age, WRAT, PMAT, and PVRT scores are reported as mean ± standard deviation.

### B. Data Preprocessing

All MRI data were acquired on a 3T Siemens TIM Trio scanner. Functional images were preprocessed using SPM12, following standard steps including motion correction, co-registration, normalization to MNI space, and spatial smoothing with a 3 mm FWHM Gaussian kernel. Nuisance regressors (6 rigid-body motion parameters and mean framewise displacement) were regressed out, and the time series were band-pass filtered (0.01–0.1 Hz).

Based on the widely used Power-264 functional atlas [38], we parcellated the brain into 264 regions of interest (ROIs). The BOLD signal for each ROI was averaged within a 5 mm sphere centered at its coordinates. Following established approaches in prior literature [15], [16], these ROIs were subsequently categorized into canonical functional networks. Specifically, 264 ROIs were assigned to 13 predefined functional brain networks: the visual network (VIS), default mode network (DMN), cinguloopercular network (COP), frontoparietal network (FPT), dorsal attention network (DAT), ventral attention network (VAT), auditory network (AUD), salience network (SAL), subcortical network (SBC), cerebellar network (CB), memory network (MEM), somatomotor hand network (SMT-hand), somatomotor mouth network (SMT-mouth) and uncertain network (UNK). For each participant and each fMRI paradigm, we computed a 264×264 symmetric FC matrix using Pearson correlation between the time series of all ROI pairs.

The diagonal elements (self-connections) were excluded, and only the upper triangular entries were retained, resulting in 34,716 unique FC features per subject per session, given by *n*(*n*−1)*/*2 where *n* = 264. These values were then transformed using Fisher’s *z*-transformation to stabilize variance and improve the normality of residuals, facilitating valid inference under the LMM framework.

To focus on the most robust FCs, we selected the top 5% of connections ranked by absolute FC strength (1,736 connections) within each paradigm. We then took the intersection of these top connections across all three paradigms, retaining only those that consistently ranked in the top 5% throughout. The top 5% threshold was selected in line with prior functional connectivity studies that retain a small proportion of the strongest edges to reduce noise and generate a sparse the network (e.g., use of top 10% in language network studies [39]). In our case, we empirically set the threshold to the top 5% of connections, which provided a stable representation of core functional relationships across paradigms without introducing excessive sparsity in the network. The retained FCs were mapped to canonical functional networks for visualization and used as features in subsequent predictive modeling.

### C. Framework Overview

The goal of our work is to characterize how demographic and experimental factors jointly impact FC, while accounting for individual variability across subjects. Each participant completed three fMRI sessions yielding multi paradigm FC per subject. This repeated-measures structure introduces dependencies among observations from the same individual, motivating a hierarchical modeling strategy that distinguishes population-level effects from subject-specific deviations.

For subjects *i* (*i* = 1, …, *q*), each has three paradigms *j* (*j* = 1, 2, 3), and a symmetric 264×264 FC matrix **F**_*i,j*_. Each element **F**_*i,j*_(*r, s*) represents the Pearson correlation between the ROI *r* and ROI *s*. To analyze connectivity patterns at the level of individual ROI pairs, we treated each pair of FC (*r, s*) as a separate dependent variable. Specifically, for each FC edge (*r, s*), we concatenated its Fisher-transformed values across all subjects and paradigms to form a response vector:

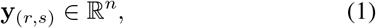

where *n* denotes the number of observations across all subjects and paradigms. Each **y**_(*r,s*)_ thus captures how the strength of the same functional connection varies across individuals and paradigms. In the following, we omit the index of (*r, s*) for simple modeling.

The design matrix of fixed effects, **X** ∈ℝ^*n*×*p*^, where *p* denotes the number of fixed-effect parameters, includes population level information such as age, sex, and paradigm to capture systematic demographic and task-related effects on FC. The design matrix of random effects, **Z** ∈ℝ^*n*×*q*^, encodes subject identity by mapping each of the *n* observations to its corresponding subject among the *q* unique participants, thereby associating each observation with a unique participant for the random intercept. The corresponding random-effect coefficients, **b** ∈ℝ^*q*^, represent subject-specific deviations in FC strength from the population mean, reflecting reproducible individual “fingerprints” of brain organization. Finally, ***ε*** ∈ℝ^*n*^ represents residual variability due to measurement noise and paradigm-specific fluctuations.

In summary, FC served as the dependent variable (**y**), with fixed effects (**X**) capturing shared, population-level influences, and random effects (**Zb**) modeling stable within-subject structure. This hierarchical formulation provided a principled foundation for both the classical LMM and its nonlinear neural extension NMM, which was employed to capture higher-order dependencies in the data.

### D. Classical LMM Revisit

Building on the hierarchical structure outlined earlier, we employed the LMM as a baseline analytical framework to assess how fixed and random effects collectively account for FC. The LMM improves upon standard linear regression by simultaneously modeling both population-level fixed effects and subject-specific random effects. This approach effectively captures the repeated-measures and nested nature of fMRI data, offering a more appropriate representation of its inherent dependencies. Formally, the model is expressed as:

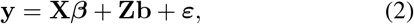

where ***β*** are the fixed-effect coefficients and **b** are the random-effect coefficients.

The random effects are assumed to follow a multivariate normal distribution,

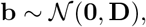

where **D**∈**ℝ**^*q*×*q*^ is a positive semi-definite covariance matrix of variance components. Here we use **D**(***θ***) to represent **D**. Residual errors are assumed to be independent and normally distributed,

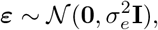

where **I** is the *n* × *n* identity matrix; **b** and ***ε*** are independent.

### E. NMM

While linear mixed-effects models combine fixed and random effects, their reliance on simple linear relationships often fails to capture the complexity of real-world data. The NMM framework [40], [41] offers various extensions that can incorporate nonlinearities in both fixed and random effects. However, in this work we focus on a specific implementation that replaces only the linear fixed-effects term with neural networks while preserving the linear hierarchical structure of random effects, as this approach best suits our data characteristics and the random-intercepts design.

Formally, we formulate our proposed NMM as:

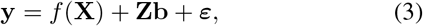

where *f* (·) denotes a multi-layer neural network. In practice, *f* (·) can be any differentiable neural architecture (e.g., multilayer perceptron (MLP)). In our work, we adopt MLP to capture the nonlinearity of the fixed effects.

Given that our data involve a single random effects categorical variable with *q* levels, the random intercepts model provides the appropriate structure. All subsequent developments are therefore derived under this specific modeling framework.

We assume a standard random-intercepts structure in which

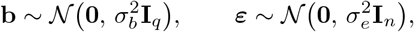

with 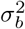 the between-subject variance and 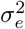 the residual (within-subject) variance. Let

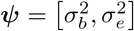

collect the variance components. Under these assumptions, the marginal distribution of **y** is

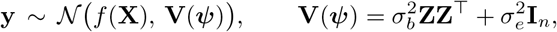

where **Z** ∈ℝ^*n*×*q*^ maps observations to subjects. Given that our data involve a single categorical random-effect factor (individual variance) with *q* levels, this random-intercepts model provides a natural and parsimonious covariance structure.

Building on likelihood-based formulations of mixed models and their neural extensions [40], [41], training proceeds by minimizing the negative log-likelihood (NLL) of the marginal distribution:

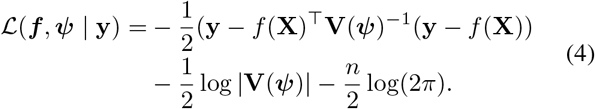

#### Model Training

We optimize this objective jointly over both the neural network parameters *f* (·) and the variance components *Ψ*.

To enable end-to-end training, the NLL loss in Eq. (5) is implemented as a differentiable objective that supports automatic differentiation. At each training epoch, the NLL is evaluated on small mini-batches (typically 30–50 observations), and gradients are propagated through both the neural and variance parameters using mini-batch stochastic optimization with Adam. The variance terms are constrained to be positive by optimizing their logarithms, which improves numerical stability.

When the covariance matrix **V** has a tractable form, the derivatives of the NLL can also be computed explicitly as follows:

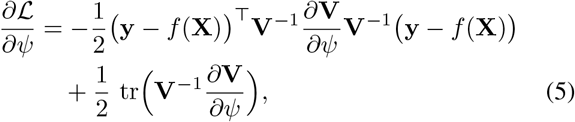

and

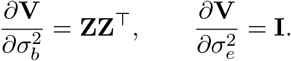

For random-intercepts models where **V** is block-diagonal, both the inverse and the log-determinant of **V** decompose across clusters, allowing the NLL and its gradient to be computed efficiently within each mini-batch.

This formulation makes it possible to integrate the hierarchical structure of linear mixed models with the representational flexibility of deep neural networks, trained jointly under a principled likelihood-based objective.

After training, the random effects are estimated using the best linear unbiased predictor (BLUP):

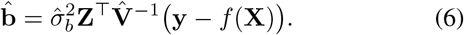

For the random-intercepts case, the closed-form solution for cluster *j* with *n*_*j*_ observations is:

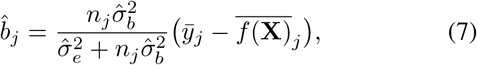

where 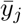 and 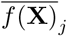 denote the sample and predicted means for cluster *j*, respectively.

#### Model Inference

For prediction, if test subjects were included during training, both fixed and random components are used. Let 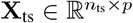 denote the fixed-effects design matrix for the test samples, where *n*_ts_ is the number of test observations and *p* is the number of fixed-effect parameters. Similarly, let 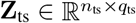 represent the random-effects design matrix for the test data, where *q*_ts_ is the number of subjects. The prediction is then given by:

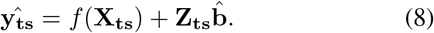

#### Model Interpretation in the FC Context

In the proposed NMM framework, each term maps directly to a component of the FC study design. The vector **y** contains the Fisher-transformed correlation values for a specific pair of FC across all subjects and paradigms. The fixed-effects component *f* (**X**) models the combined influence of age, sex, and paradigm on FC strength, allowing for non-linear and interactive effects. This is consistent with prior work modeling developmental changes in FC or related metrics with age effects and sex covariates [42], [43], as well as task-based studies where FC is explicitly modeled as a function of task context and its interactions with demographic factors [44]. In addition, another study has demonstrated that age and sex jointly shape cortical dynamics via significant age-by-sex interaction effects [45], reinforcing the need to allow for interactive fixed effects in *f* (**X**). The random-effects term **Zb** accounts for consistent individual differences across scanning sessions, capturing the subject-specific variability in FC that is not explained by the fixed factors. The residual term ***ε*** represents unexplained variation, including measurement noise and transient fluctuations unique to each scan. This model structure aligns with the repeated-measures nature of the data, separating population-level trends from stable individual differences.

#### Advantages over Classical LMM

The key improvement of NMM over the standard linear mixed model lies in its ability to capture non-linear and interaction effects among fixed predictors. In conventional LMMs, the relationship between covariates and the outcome is constrained to be linear and additive. In contrast, the neural network component *f* (·) can learn more complex functional forms from the data, which may better reflect the actual relationships between demographic variables, task conditions, and functional connectivity. This is particularly relevant for FC data, where the combined effects of age, sex, and paradigm may not adhere to a simple linear model. By incorporating a flexible non-linear fixed-effects component while retaining the linear random-effects structure, NMM provides a more adaptive framework for modeling FC, as demonstrated in recent methodological work integrating neural networks with LMMs [40], [41].

## III. Experimental Results

### A. Robust FCs Shared Across Paradigms

Figure 2 summarizes the FC selection procedure. Panels A–C show, for emotion, nback and rest tasks respectively, the heatmaps of the top 5% FCs (colored), with the remaining 95% left blank. Panel D displays the intersection of these selections across paradigms, yielding 1,243 consensus FCs (≈3.6% of all FCs).

**Fig. 2.**
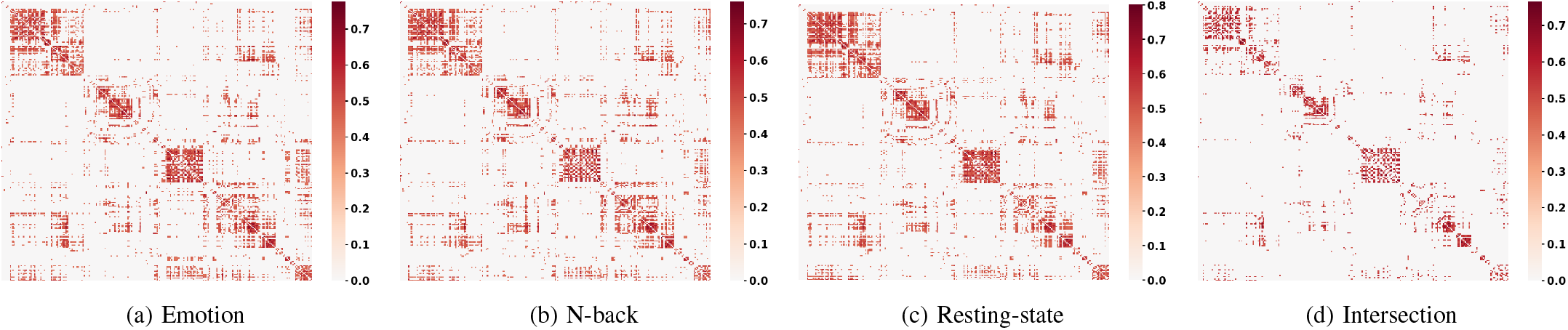
Extracted FCs for each paradigm and their intersection.

To contextualize these features, we mapped the 1,243 FCs to canonical functional networks and visualized their distribution using a circos plot (Figure 3). This view highlights the relative contributions of intra and inter network connections among the retained FCs, providing a compact summary of the network architecture represented in the final feature set.

**Fig. 3.**
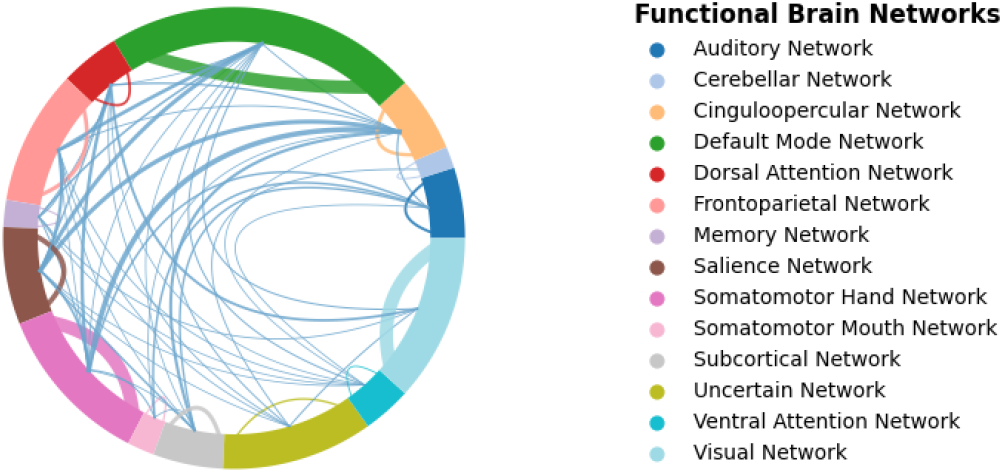
Selected FCs map onto intra- and inter-network connections across canonical brain networks. Arc length is proportional to the number of ROIs in each network, and line width is proportional to the number of functional connections. Intra-network connections are shown in the color of the corresponding network, while inter-network connections are shown in blue-gray.

### B. Goodness of fit and Predictive Performance

In this study, the dataset was randomly divided into training and testing subsets, with 80% of the data used for model training and 20% reserved for testing. The NMM model was implemented in `TensorFlow/Keras` and trained using the Adam optimizer. The architecture consisted of two hidden layers with a rectified linear unit (ReLU) activation and fully connected units of 10 and 5 neurons, respectively. The model estimated population-level fixed effects and subject-specific random effects. Training and testing were performed iteratively across all target variables, and model performance was quantified using the mean squared error (MSE) between predicted and observed FC values in the test set. For baseline comparison, classical LMMs were fitted using the `statsmodels` package, with the same fixed and random-effect structure as the neural implementation. Model fit was assessed by MSE as well.

Although only a single random 80/20 train–test split was reported, preliminary analyses indicated that alternative splits (e.g., 70/30 or 90/10) produced highly similar MSE values. The 80/20 proportion was therefore adopted as a standard and widely accepted compromise between training-set sufficiency and test-set reliability in neuroimaging and deep-learning research [46], [47]. Similarly, minor variations in the network size (e.g., 8–4 or 12–6 neurons per layer) yielded nearly identical predictive performance, consistent with the common recommendation to favor minimal architectures that are sufficient for convergence without over-parameterization [48], [49]. Given that the main purpose of this work was to compare the mixed-effects formulation between the classical and neural implementations rather than to exhaustively optimize hyper-parameters, and that repeated 100-fold resampling would considerably increase computational cost due to the multi-output random-effects structure, we used a single representative 80/20 split for reporting. This design ensured methodological consistency and computational efficiency while maintaining generalizable model evaluation.

For each of the 1,243 selected FCs, we trained NMM and classical LMM using fixed effects (age, sex, paradigm) with a subject-level random effect. We computed the mean test set MSE across all 1,243 FCs. NMM showed lower error than the classical LMM (0.0258 ± 0.0044 vs. 0.0359 ± 0.0064; mean ± SD), as shown in Figure 4.

**Fig. 4.**
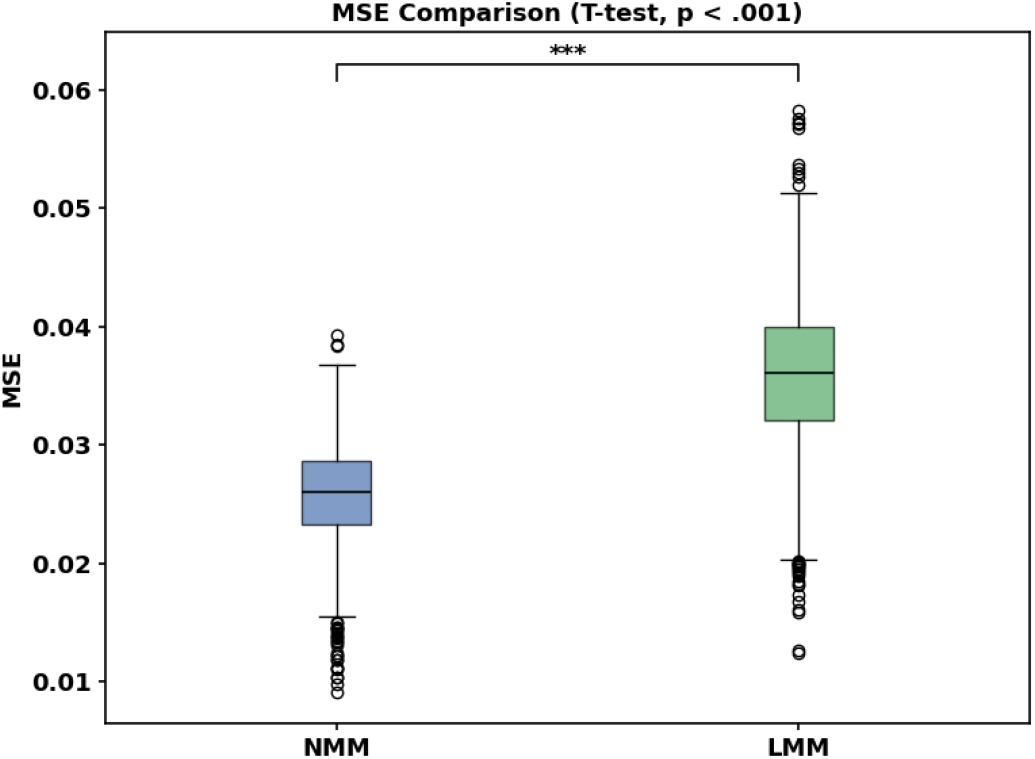
Mean MSE comparison between NMM and LMM across 1,243 FCs on the test set

Additionally, we visualized a randomly chosen FC by plotting predicted–observed scatter for NMM and LMM, as shown in Figure 5. The Pearson correlations are *r* = 0.5214 (*p <* .001) for NMM and *r* = 0.2315 (*p <* .001) for LMM. All metrics were computed on test data, and the single FC behavior matches the aggregate MSE advantage.

**Fig. 5.**
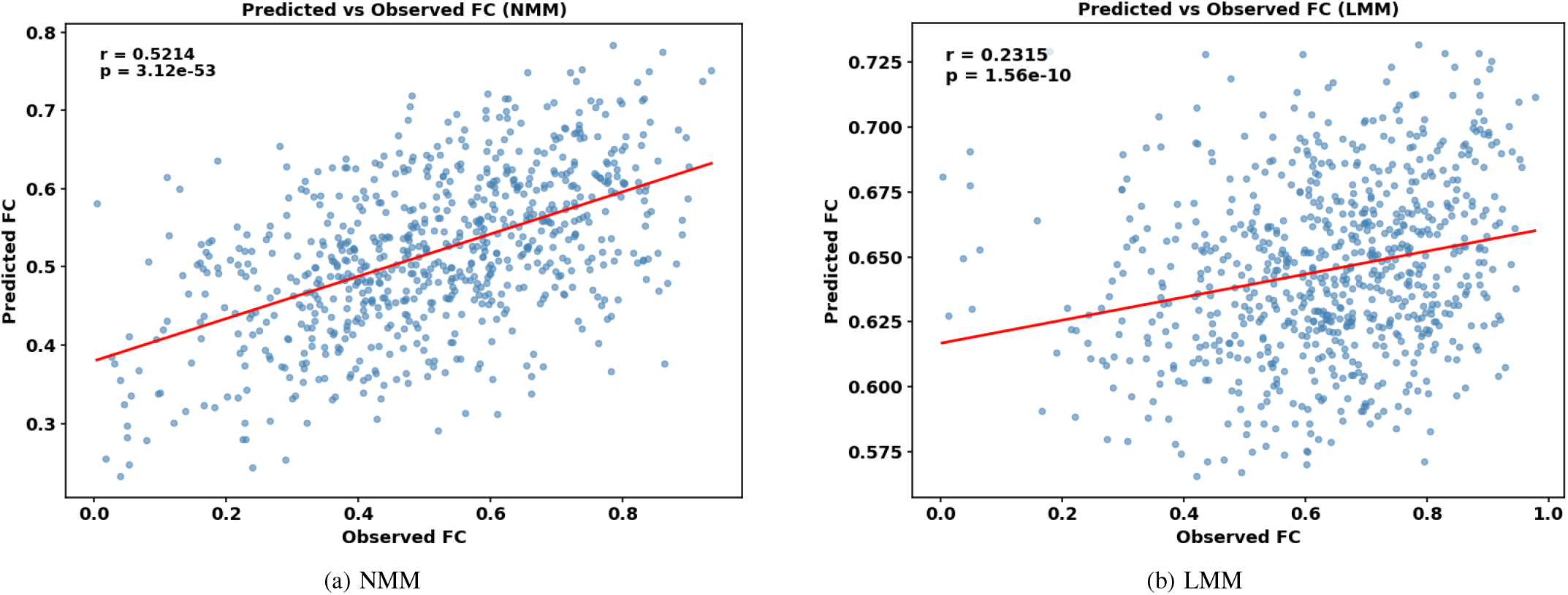
Predicted versus observed FC values for a representative connection under NMM and LMM.

### C. Interpreting Fixed Effects Using SHAP

To elucidate the mechanistic contributions of fixed effects within our NMM framework, we employed SHAP analysis, focusing specifically on the fixed effects component while excluding random effects. The fixed effects subnetwork was isolated, and SHAP values were computed using KernelExplainer with 200 randomly sampled background observations to ensure computational efficiency.

At the global level as shown in Figure 6, mean absolute SHAP values revealed that paradigm contributed most substantially to fixed effects predictions, followed by age and sex variables. This hierarchical importance was consistent across multiple functional connectivity targets, suggesting robust feature contribution patterns. To localize these effects within specific functional brain networks, we aggregated the 1,243 FCs according to their corresponding brain networks and computed mean absolute SHAP values for each network pair. This analysis distinguished between intra network (diagonal elements) and inter network (off-diagonal elements) connectivity contributions. Panels (a)–(c) display the maps for age, sex, and paradigm, respectively. Notably, sex demonstrated uniformly low SHAP contributions across all network pairs, aligning with its minimal global importance. In contrast, paradigm exhibited concentrated high contributions within specific network interactions, particularly between DMN and DAT, DMN and SBC, SMT-hand and SBC, and SMT-mouth and COP. Age-related contributions displayed a distinct pattern, with strongest effects observed in AUD-SBC, COP-DMN, DMN-DAT, VAT-DAT, and SMT-mouth-SBC.

**Fig. 6.**
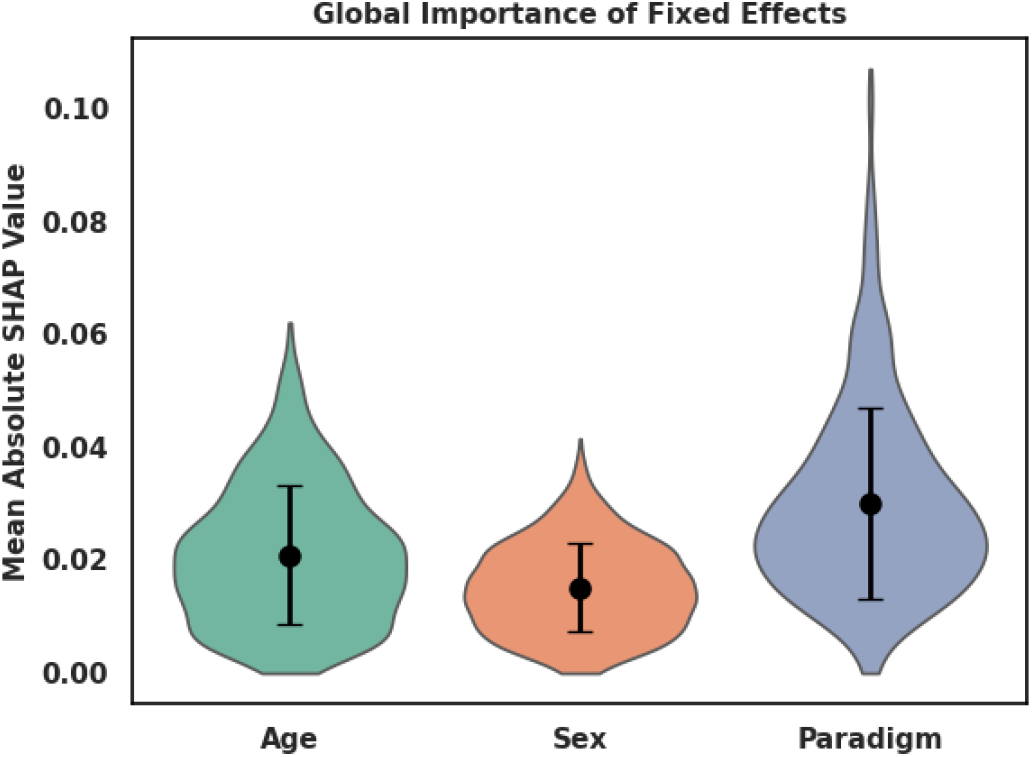
Global importance of fixed effects.

These network-specific SHAP patterns provide mechanistic insights into how demographic and experimental factors differentially modulate distinct functional brain systems, with paradigm primarily influencing cognitive control networks, while age effects span sensory, attention, and default mode systems.

### D. Random effects Variability

Subject level random effects are assumed to follow a Gaussian distribution with mean zero. We therefore focused our analysis on the variance components to characterize between-subject variability in FC patterns. The magnitude of this variance reflects the degree to which individual subjects deviate from the population average, with larger values indicating more pronounced individual “fingerprints” within specific functional systems.

For each FC, we computed the standard deviation of subject random intercepts across participants. We then grouped FCs by brain networks and summarized intra networks by their network level mean and dispersion of random effect variance. Intra brain networks were assessed using two sample t-tests. Figure 8 reports the distribution of subject variance across networks. The analysis reveals substantial heterogeneity in individual variability patterns across different brain systems. The VAT demonstrates significantly elevated random effect variance compared to other systems (*p <* .001), suggesting particularly strong individual differentiation in attention-related circuits. Conversely, VIS exhibit markedly lower variance, indicating more constrained individual differences in primary sensory processing regions.

These findings align with known neurobiological principles: higher order cognitive networks supporting attention and control processes demonstrate greater individual specialization, while primary sensory systems maintain more conserved organizational patterns across individuals [50]–[52]. The robust statistical separation between networks underscores the utility of mixed effects modeling for capturing meaningful individual differences in functional brain organization.

### E. Cognitive Associations of Individual Variability

We used the subject level random effects estimated by the NMM as individual features and investigated whether individual level random effects could predict standardized cognitive assessments. For each participant, we concatenated the random effects estimates across the 1,243 FCs to form a comprehensive individual neural profile. These subject vectors served as input features for predicting three distinct cognitive scores: WRAT, PMAT and PVRT.

We employed a straightforward MLP to establish the mapping between individual differences and cognitive performance. The network architecture, implemented through scikitlearn’s MLPRegressor, consisted of a single hidden layer with 100 neurons using ReLU activation functions, optimized via Adam solver with default learning rate. The dataset was partitioned using an 80/20 train-test split with random state stabilization (random state=42) to ensure reproducible evaluation. Model performance was quantified by computing Pearson correlation coefficients between predicted and observed scores in the held-out test set, with statistical significance assessed using two-tailed p-values.

The results demonstrate that individual neural variability patterns captured by random effects contain meaningful information about cognitive functioning. For WRAT, the model achieved a correlation of *r* = 0.2741 (*p <* .01) between predicted and observed scores. PMAT showed a predictive correlation of *r* = 0.2454 (*p <* .01), while PVRT exhibited the strongest association at *r* = 0.3369 (*p <* .001).

Figure 9 illustrates the test set predictions for each cognitive measure, revealing the strength and consistency of these relationships across individuals. The scatter plots show clear alignment along the identity line, particularly for PVRT, indicating that our model successfully captures systematic individual differences in cognitive performance from neural random effects alone.

Notably, the varying predictive accuracy across cognitive domains suggests differential neural representation of these abilities. The stronger prediction of PVRT implies that this cognitive domain may have more distributed neural substrates that are effectively captured by our whole-brain connectivity approach. In contrast, the more modest prediction of PMAT may reflect either more localized neural representation or greater measurement variability in this domain.

## IV. Discussion

This study introduced a novel approach which combines the statistical model–LMM with popular deep learning framework for analyzing multi-paradigm FC. Unlike previous studies, NMM emphasizes individual variability and captures nonlinear relationships, leading to improved model fit of FC. The proposed method effectively handles the complex, hierarchical structure of neuroimaging data. Our results show that the NMM not only achieves a better fit than classical LMM but also produces highly interpretable findings. Crucially, we demonstrated that the model’s random effects components directly parameterize stable trait-level biomarkers, accurately predicting standardized intelligence scores.

### Why nonlinearity from NMM?

The consistently superior predictive performance of NMM across 1,243 FCs underscores its advantage over the classical LMM. This advantage stems from the model’s capacity to capture the complex, non-linear relationships among demographic variables (e.g., age, sex), task paradigms, and FC that are ubiquitous in fMRI data. Traditional LMMs, constrained by their linear assumptions, often fail to account for the complexities inherent in real-world data [53], [54]. For instance, the relationship between age and FC is rarely constant across the developmental lifespan. The neural network component of NMM flexibly adapts to these non-linearities and higher-order interactions (e.g., age-by-paradigm), leading to more accurate predictions.

Furthermore, the NMM is exceptionally well-suited to the nested structure of multi-paradigm fMRI studies, where repeated measurements are collected from the same individual. This design aligns with the concept of the functional connectome as a unique subject-specific fingerprint, as demonstrated by Finn et al. [16]. The biological relevance of the individual variability captured by our model is further supported by its predictive utility. Cai et al. have shown that individual differences in FC can predict cognitive behavior [15]. In line with this, the random effects estimated by our NMM successfully predicted cognitive performance, indicating that the captured subject-level variance represents meaningful biological signal rather than mere noise. By integrating the flexibility of neural networks with the structured framework of mixed models, our hybrid NMM offers a powerful and practical tool for the field. It enables the modeling of large-scale FC from a modest set of covariates while retaining a clear random-effects structure and supporting post-hoc interpretation of the fixed effects via SHAP, all within a statistically rigorous framework.

### Specific functional networks and individual variability

The SHAP maps and the random effects variance point to a coherent network story. First, paradigm effects are broad and cross-network. In the inter + intra matrices (Fig. 7), high SHAP for paradigm concentrates in DMN and DAT, DMN and SBC, SMT-hand and SBC, and SMT-mouth and COP. This pattern is consistent with the idea that task context modulates large-scale control and SMT–SBC pathways supporting performance. Previous studies have demonstrated that task engagement dynamically reconfigures cortico–subcortical circuits to optimize behavioral performance, with task-induced changes in corticostriatal and thalamocortical connectivity mediating cognitive control and working-memory processes [55] By contrast, sex contributions are low and spatially diffuse, matching the global SHAP finding and suggesting that, after controlling for age and paradigm, sex accounts for limited fixed-effect variance in this cohort. Stronger SHAP for age appears in AUD-SBC, COP-DMN, DMN-DAT, VAT-DAT and SMT-mouth-SBC. Previous studies have demonstrated clear developmental trajectories in large-scale control and attention systems, as well as in sensory–subcortical integration. For example, Fair et al. showed that control networks (frontoparietal and cingulo-opercular) differentiate and shift from local to more distributed architectures across development, and that default–attention couplings reorganize with age [56]; Dosenbach et al. further demonstrated that multivariate patterns of functional connectivity predict individual brain maturity from childhood to adulthood [57]. Building on cortico–subcortical pathways, these loci align with systems where developmental trajectories in control, attention, and sensory–subcortical integration have been reported, providing a network-resolved view of how age enters the fixed component.

**Fig. 7.**
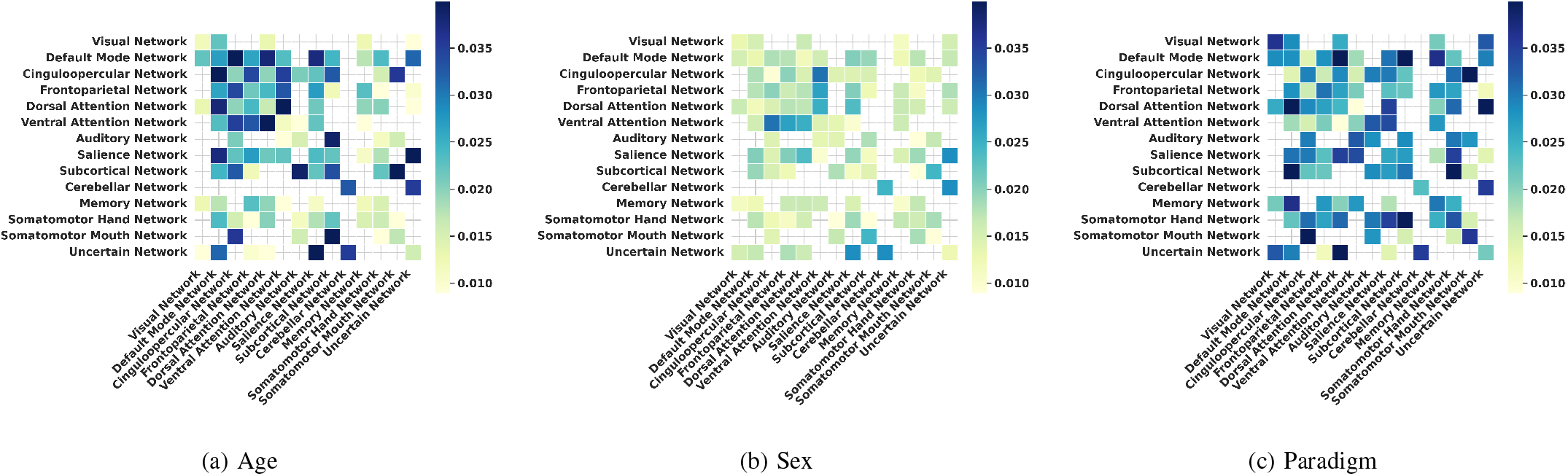
SHAP values for age, sex and paradigm in Functional Network Connectivity.

**Fig. 8.**
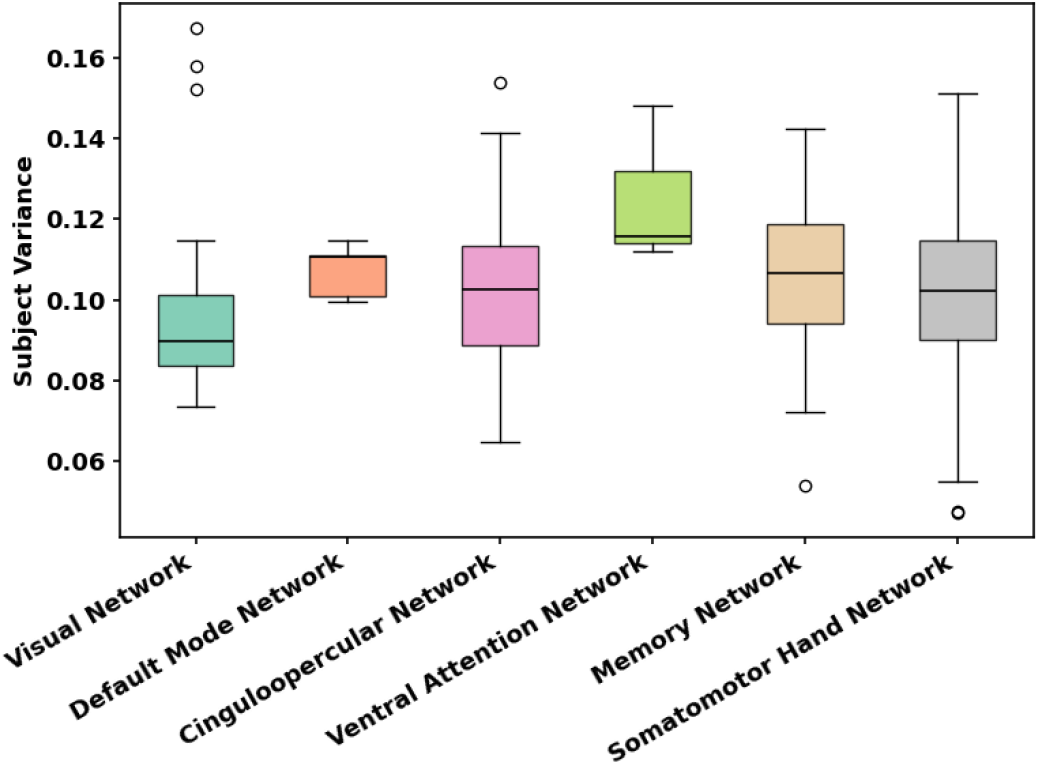
The random effects of subjects across different intra-networks

**Fig. 9.**
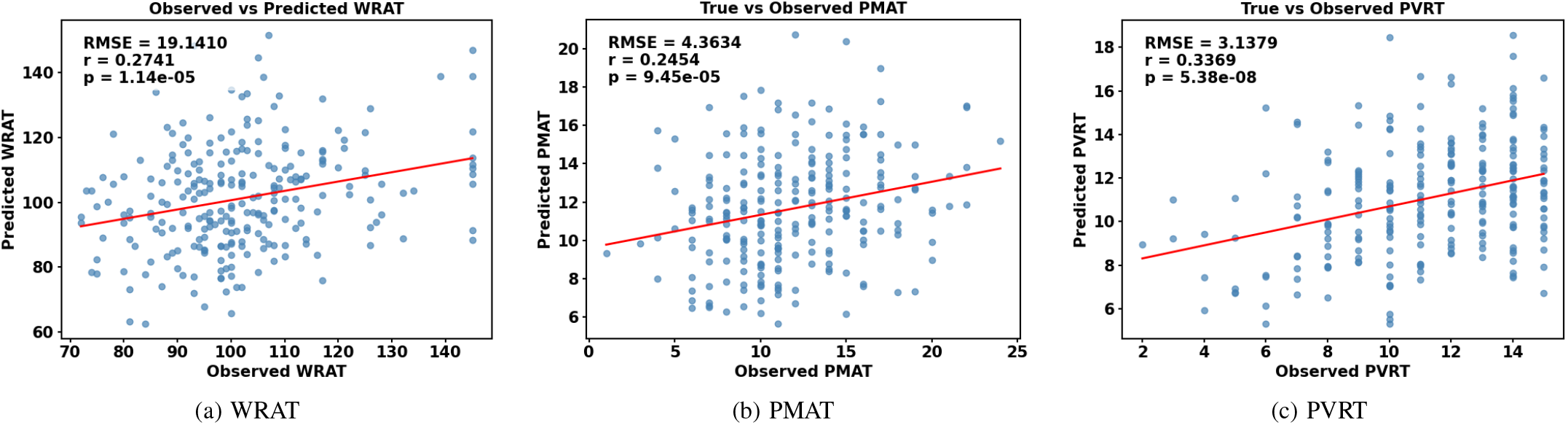
The association between the extracted FC features and cognitive behavioral measures.

Second, the random effects variance are most significant where subject specificity is strongest. The boxplots in Fig. 8 (random-effects variance by network) show significantly higher subject variance in VAT and lower variance in VIS. This pattern aligns with prior evidence that heteromodal association cortices—particularly attention and control networks—exhibit the greatest inter-individual variability, whereas unimodal sensory areas such as the visual cortex are markedly more homogeneous across people [50], [58], [59]. VAT supports reorienting and salience detection, functions known to vary across individuals and tasks; high variance here suggests that the NMM random part absorbs meaningful idiosyncratic engagement of attentional control. VIS, in contrast, shows constrained variability, consistent with the stability of primary sensory systems [60]. DMN and COP fall between these extremes, with moderate variance and many points near the median, reflecting a mix of stable baseline organization and task-related modulation. Together, these findings reinforce that the individual fingerprint is not uniformly distributed across systems but is concentrated in higher-order control circuits.

Taken together, the SHAP and variance views are complementary: SHAP localizes where the fixed covariances drive predictions (paradigm broadly; age in targeted network pairs; sex weakly across the connectome), while random-effects variance indicates where subject-specific departures from the population mean are largest (VAT) or smallest (VIS). This division of labor is desirable: the fixed component captures shared, covariant-linked structure, and the random component captures stable individual signatures. The coherence between these two lenses increases the confidence that NMM is modeling signal rather than noise.

## V. Conclusion

In this paper, we introduced NMM to capture the relationship between FC and simple covariates—age, sex, and paradigm while retaining a subject level random effects. Across selected FCs, the nonlinear fixed component delivered consistently lower MSE than a classical LMM. SHAP analyses clarified that fMRI paradigm drives most of the fixed effect signal, whereas the random effects variance revealed robust individual “fingerprints” of fMRI, which also carried predictive value for cognitive scores. Taken together, our findings show that bringing neural networks into a mixed effects formulation can better fit FC data while maintaining a transparent random-effects structure and supporting post-hoc explanations of the nonlinear fixed effects. The results align with prior reports on individual variability and task modulation in fMRI, and this framework offers a practical route for modeling large scale FC with modest inputs.

### Limitations and Future Work

Our inputs are intentionally simple (age, sex, paradigm) and we use individual variation as random effects; more comprehensive model specifications could add random slopes for family structure, or site effects, and extend to dynamic FC. We selected robust FCs via a multi-paradigm intersection; future work could test sensitivity to other feature criteria and to alternative atlases or parcellation schemes. Finally, external validation across cohorts will clarify generalization. Despite these limits, the present results show that a nonlinear mixed-effects approach can recover meaningful population effects and individual fingerprints from modest covariates, while providing a structured basis for post-hoc interpretation.

## Acknowledgment

This work was supported in part by National Institutes of Health (NIH), USA under Grants R01 EB036247, R01 GM109068, R01 MH104680, R01 MH107354, P20 GM103472, R01 REB020407, R01 EB006841, P20-GM144641 and in part The U.S. National Science Foundation (NSF) under Grant 1539067.

## Notes

This work was partially supported by NIH Grants R01 EB036247, R01 GM109068, R01 MH104680, R01 MH107354, P20 GM103472, R01 REB020407, R01 EB006841, P20-GM144641 and NSF Grant #1539067.

### Competing Interest Statement

The authors have declared no competing interest.

